# Demographic pathways linking sex-chromosome system to adult sex ratio variation across tetrapods

**DOI:** 10.1101/2024.11.17.624031

**Authors:** Ivett Pipoly, Veronika Bókony, Jean-Michel Gaillard, Jean-François Lemaître, Tamás Székely, András Liker

**Affiliations:** HUN-REN-PE Evolutionary Ecology Research Group, University of Pannonia, H-8210 Veszprém, Pf. 1158, Hungary; Behavioral Ecology Research Group, Center for Natural Sciences, Faculty of Engineering, University of Pannonia, Pf. 1158., H-8210 Veszprém, Hungary; Department of Evolutionary Ecology, Plant Protection Institute, HUN-REN Centre for Agricultural Research, Herman Ottó út 15, H-1022 Budapest, Hungary; Université Lyon, CNRS, Laboratoire de Biométrie et Biologie Évolutive, France; HUN-REN-DE Reproductive Strategies Research Group, University of Debrecen, H-4032 Debrecen, Hungary; Milner Centre for Evolution, Department of Life Sciences, University of Bath, Bath BA2 7AY, United Kingdom; Debrecen Biodiversity Centre, University of Debrecen, H-4032 Debrecen, Hungary

**Keywords:** sex chromosomes, birth sex ratio, sex-specific mortality, maturation, time of sexual maturity, phylogenetic comparative analyses, tetrapods

## Abstract

Sex chromosomes determine male and female phenotypes, and the resulting sex differences can have significant impacts on ecology and life history. One manifestation of this link is that ZZ/ZW sex-determination systems are associated with more male-skewed adult sex ratio (ASR, proportion of males in the adult population) than XY/XX systems across tetrapods (amphibians, reptiles, birds, and mammals). Here we investigate four demographic processes: male and female offspring production, sex differences in juvenile and adult mortalities and in timing of maturation that can contribute to ASR variation between XY/XX and ZZ/ZW systems, using phylogenetic analyses of a large dataset collected from tetrapod species in the wild. We show that sex differences in adult mortality reliably predict ASR, and it is also more male-biased in XY/XX species than in ZZ/ZW species. Sex differences in juvenile mortality or in maturation time also contribute to ASR skews, but do not differ consistently between XY/XX and ZZ/ZW systems. Phylogenetic path analyses confirm an influence of sex-determination system on ASR through sex-biased adult mortality. Thus, these results infer that sex chromosomes can impact, via demographic pathways, frequency-dependent selection emerging from the relative number of males and females. We call for follow-up studies to uncover the potentially complex web of associations between sex determination, population dynamics, and social behaviour.

## Introduction

A growing body of evidence suggests that adult sex ratios (ASR, the proportion of males in the adult population) are often skewed in animal populations[1–4]. ASR is a central demographic property of populations because it plays a key role in population dynamics, for example through its effects on reproductive rates and population viability[5,6]. ASR also affects the number of competitors for mating and the number of mates available to sexually mature adults in the population[7–9], thus it is associated with variation in reproductive behaviour including mating system, parental care, divorce rate and extra-pair paternity[10–14]. ASR is also associated with variations in life-history traits such as age at maturation[15], and various indicators of sexual selection including sexual dimorphism in body size and/or ornamentation[16,17]. However, how skewed ASRs emerge in the wild remains poorly understood.

Sex chromosomes influence male and female phenotypes in many organisms, and recent studies have uncovered that the genetic sex-determination system (GSD, defined by the type of sex chromosomes) may lead to sex differences in demographic and life-history traits. For example, higher mortality and/or ageing rates have been reported in the heterogametic sex (i.e. XY males in species with XY/XX sex-chromosome systems, and ZW females in species with ZZ/ZW systems) compared to the homogametic sex (i.e. XX females and ZZ males, respectively) across dioecious animals[18–20] and plants[21]. Phylogenetic analyses also suggest that the type of GSD may also influence ASR variation among species since ASR was more male-skewed in ZW species (i.e. species with ZZ/ZW systems) than in XY species (i.e. species with XY/XX systems). Although it is critical to understand how sex-chromosome systems contribute to the observed inter-specific variability in ASR, the demographic pathways by which ASR is linked to the type of GSD and whether these pathways differ between GSD types have never been investigated so far. There are several pathways that may potentially generate a link between the type of GSD and ASR (Fig. 1). A skewed ASR can result from sex-biased offspring production. For example, this may occur due to meiotic drive linked to sex chromosomes, which can produce gametes biased towards the sex that carries the driving sex chromosome[23]. This mechanism occurs in species with either XY or ZW systems and causes biases in offspring sex ratio[24,25]. A comparative study did not support this pathway in birds (all having ZW system) because offspring sex ratio was not related to variation in ASR across species[26]. However, whether offspring sex ratios differ between XY and ZW species and whether it is related to ASR in a broader range of taxa has not been investigated yet.

**Fig. 1.**
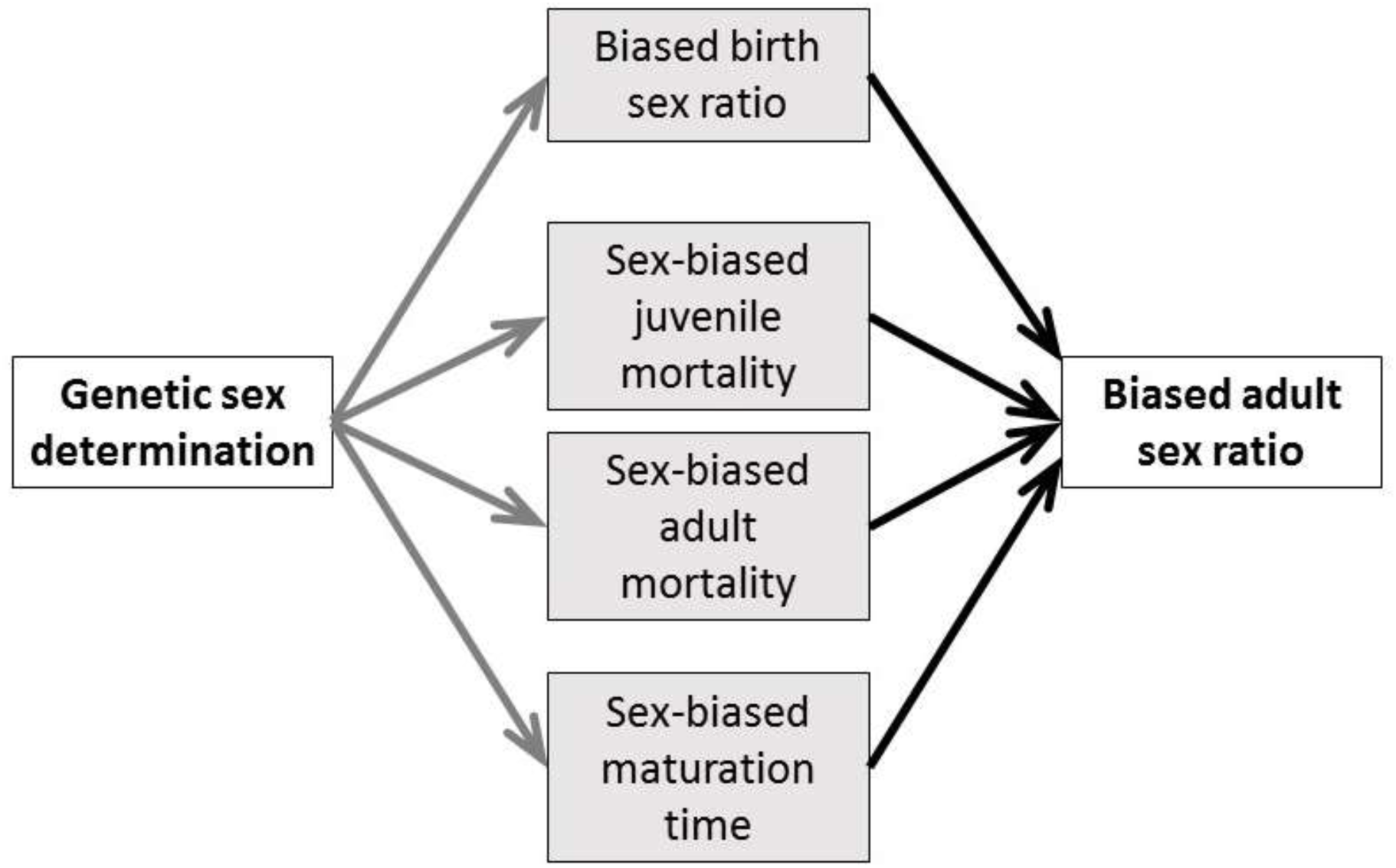
Biological and demographic pathways tested in this study through which the type of genetic sex-determination systems may influence adult sex ratio. We test (i) which of the demographic variables differ between ZZ/ZW and XY/XX sex-determination systems (grey arrows), and (ii) whether the demographic traits predict adult sex ratio (black arrows). Also, we (iii) carry out a phylogenetic confirmatory path analysis that compares the fit of path models that contain different sets of relationships (grey and black arrows together).

GSD may also be associated with ASR through its effects on sex-specific mortality at the juvenile and/or adult life stages (Fig. 1). Sex chromosomes can influence sex-biased mortality through several mechanisms (reviewed in[27]) for example due to the more frequent expression of harmful recessive mutations in the heterogametic sex (unguarded sex chromosome hypothesis[1,28]), or by the accumulation of deleterious mutations and repetitive DNA elements on the Y (or W) chromosome (the toxic Y hypothesis[29]). In line with these hypotheses, shorter lifespan and lower survival of the heterogametic sex have been reported in various taxa (see above), and sex-biased adult mortality is the strongest predictor of skewed ASRs in birds[1,26].

Sex differences in the time of sexual maturation (called maturation bias hereafter) can also contribute to ASR variation[2,30]. Sexual selection and/or fertility selection may lead to sex differences in maturation age[15], resulting in ASRs skewed towards the sex with faster maturation. This maturation bias hypothesis (Fig. 1) is supported in birds and reptiles where females mature later than males in populations with male-skewed ASRs and *vice versa*[15,30]. Whilst there are specific studies that investigated one of these hypotheses in a limited number of taxa (see examples above), none of the previous analyses evaluated all three hypotheses across a broad phylogenetic context.

Here we use tetrapods – an unusually variable clade that include amphibians, reptiles, birds and mammals – to test which demographic pathways may mediate the link between GSD and ASR variation. Tetrapods exhibit both XY/XX and ZZ/ZW sex determination systems, display highly diverse ASRs[2,3,22], and detailed demographic data are available from a large number of species thanks to long-term population monitoring[31]. To link GSD to ASR, sex differences in the demographic traits should differ between XY and ZW species, and should be associated with ASR. By using detailed demographic data from 453 species from 29 orders and 124 families (Fig. 2) on birth sex ratio, sex biases in juvenile and adult mortality and maturation times (Fig. 1), first we test which demographic traits differ between XY and ZW species (grey arrows in Fig. 1). Second, we examine which demographic traits best account for variation in ASR across species (black arrows in Fig. 1), additionally testing whether these relationships are consistent between XY and ZW species. Lastly, we carry out confirmatory phylogenetic path analyses to compare the fit of our data to *a-priori* scenarios that link GSD and ASR via different demographic pathways (grey plus black arrows in Fig. 1). These analyses corroborate and extend our previous analyses and provide novel results by identifying the demographic processes that lead to skewed ASRs in tetrapods.

**Fig. 2.**
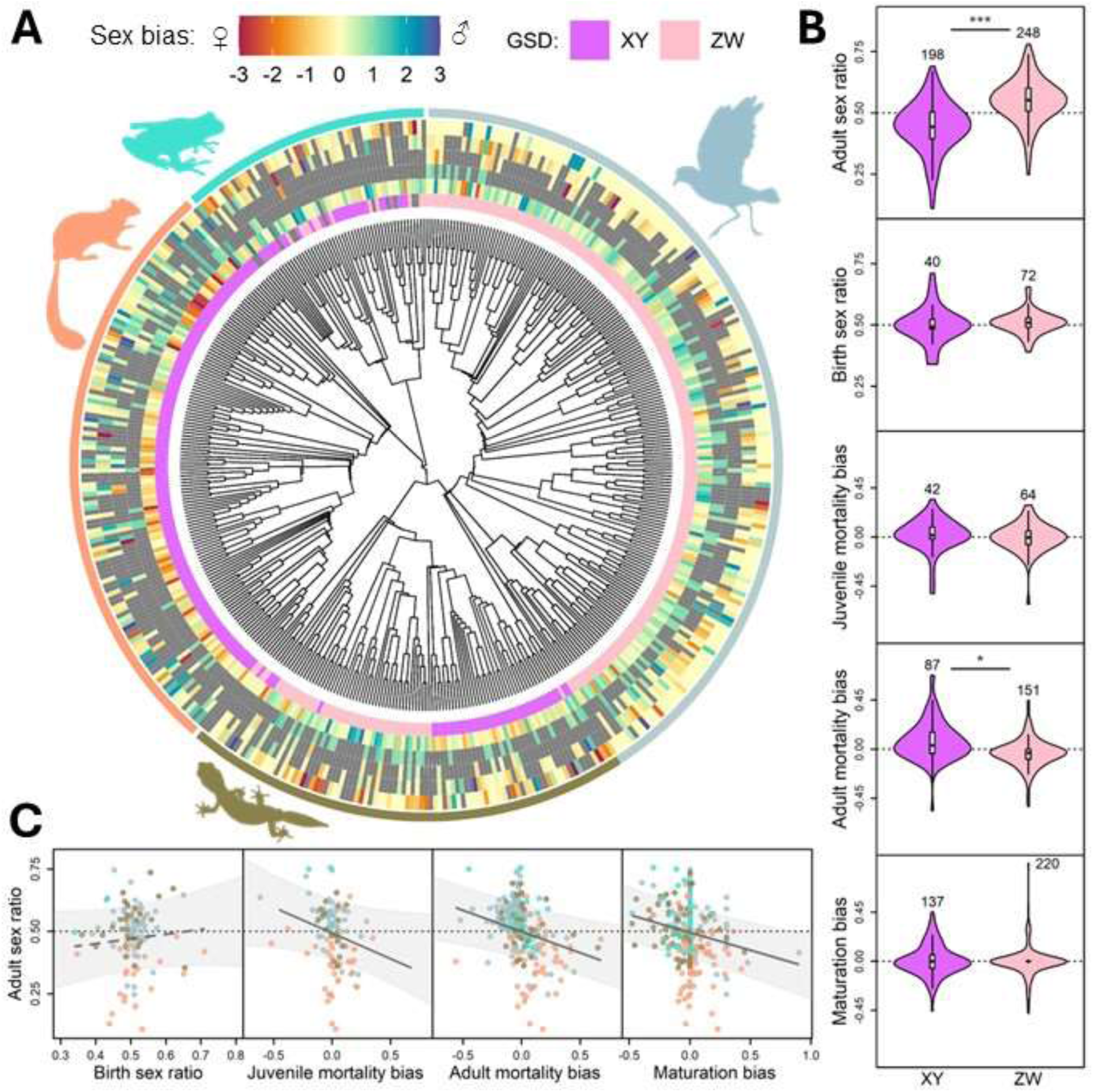
Relationships between the type of genetic sex-determination system (GSD), adult sex ratio (ASR) and sex bias in demographic traits across tetrapods. (A) Phylogenetic distribution of the six variables across 453 tetrapod species. The innermost colour circle shows the type of GSD (XX/XY and ZZ/ZW sex-chromosome systems abbreviated as XY and ZW, respectively), whereas the outer circles illustrate sex biases shown as female (red, negative values) or male (blue, positive values) bias, in the following order from inner circle to outer circle: ASR, birth sex ratio (both sex ratios depicted as deviations from 1:1), and sex differences in juvenile and adult mortality and maturation age. Bias values are plotted as standardised (z) scores centred on zero (i.e. no bias) for ease of visualisation on a comparable scale. Extreme positive and negative values are plotted as 3 and −3 to prevent outliers from obscuring the major patterns. Note that male-biased values mean male-skewed sex ratios and males having higher mortality and later maturation. Missing values are indicated by grey colour. B) Differences in ASR and other demographic traits between XY and ZW systems. In each violin plot, the black diamond and the white box represent the median and interquartile range, whiskers extend to 1.5 × interquartile range, and the polygon is a kernel density plot. Sample sizes (number of species) are shown above each plot; asterisks mark statistically significant differences (*: p < 0.05, ***: p < 0.001; see Table S2). (C) Relationships of ASR with demographic traits. Data points represent species, coloured by the four major tetrapod clades as shown by the colour of the animal silhouettes around the phylogeny (A). The regression lines are calculated from phylogenetic generalised least squares (PGLS) models (continuous lines: statistically significant relationships, p ≤ 0.01; dashed lines: non-significant relationship; see Table S2). Light grey polygons illustrate the 95% confidence bands of the regression lines assuming Brownian motion. In both (B) and (C), > 0.5 sex ratio values mean male-skewed sex ratios, while positive bias values mean higher male than female mortalities and maturation times.

## Results

### XY and ZW species exhibit different sex-dependent demographics

Species with ZZ/ZW and XY/XX sex determination systems exhibit significantly different ASRs (Fig. 2A & 2B; Table 1A & S1) consistently with our previous study that used 344 species of tetrapods[22]. ASR was a male skewed in ZZ/ZW systems and a female skewed in XY/XX systems (Table S1; Fig. 2A & 2B). The type of GSD was also associated with sex differences in adult mortality, with more male-biased mortality (i.e. higher mortality in males than in females) in XY compared to ZW species, as expected (Table 1A & S1; Fig. 2B). These effects of the sex chromosomes on ASR and adult mortality bias were medium-strong (Table 1A). However, XY and ZW species did not differ significantly in the other demographic traits, with only weak differences in birth sex ratio, juvenile mortality bias, and maturation bias (see effect sizes in Table 1A & S1; Fig. 2 A & B).

**Table 1.** Phylogenetic generalised least squares analyses of the bivariate relationships between the type of genetic sex-determination system, population demographic traits, and adult sex ratio in tetrapods. **(A)** Difference between XY and ZW species in demographic traits. **(B)** Associations between demographic traits (predictors) and the adult sex ratio (ASR, response variable). Note that the continuous variables were standardised to 0 mean and 1 SD, so the model coefficients (b) are comparable across predictors. In each model, **λ** is the phylogenetic signal (Pagel’s lambda), **R^2^** is the proportion of variance explained by the model, **n** is the number of species (number of XY species and ZW species, respectively, presented in parentheses). Statistically significant (i.e. p< 0.05) relationships are typed in bold.

| Variable | $b^* \pm SE$ | t | p | $\lambda$ | $R^2$ | n ($n_{XY} / n_{ZW}$) |
| --- | --- | --- | --- | --- | --- | --- |
| *(A) Differences between XY and ZW species in tetrapods* |  |  |  |  |  |  |
| **ASR** | 0.784 ± 0.166 | 4.715 | **< 0.001** | 0.316 | 0.048 | 446 (198/248) |
| BSR | 0.072 ± 0.201 | 0.889 | 0.376 | 0.000 | 0.007 | 112 (40/72) |
| Juvenile mortality bias | -0.202 ± 0.317 | -0.639 | 0.524 | 0.194 | 0.004 | 106 (42/64) |
| **Adult mortality bias** | -0.508 ± 0.236 | -2.152 | **0.032** | 0.307 | 0.019 | 238 (87/151) |
| Maturation bias | -0.342 ± 0.215 | -1.595 | 0.112 | 0.341 | 0.007 | 357 (137/220) |
| *(B) Associations between ASR and demographic traits in tetrapods* |  |  |  |  |  |  |
| BSR | 0.097 ± 0.093 | 1.044 | 0.298 | 0.398 | 0.009 | 112 |
| **Juvenile mortality bias** | -0.256 ± 0.098 | -2.615 | **0.010** | 0.338 | 0.061 | 108 |
| **Adult mortality bias** | -0.193 ± 0.059 | -3.251 | **0.001** | 0.288 | 0.042 | 243 |
| **Maturation bias** | -0.193 ± 0.046 | -4.203 | **< 0.001** | 0.383 | 0.046 | 365 |

\* In (A) "b" shows the difference in ASR between XY and ZW species, with XY as the reference group. In (B) "b" gives the slope of the linear regression. Because the variables are standardised to 0 mean and 1 SD, "b" values in (A) can be interpreted as Cohen's d, where values around 0.2, 0.5, and 0.8 are considered as small, medium, and large effect sizes, respectively; whereas "b" values in (B) can be interpreted as correlation effect sizes, where values around, 0.1, 0.3, and 0.5 reflect weak, medium, and strong correlations, respectively.

### Association between demographic traits and ASR

Sex bias in both juvenile and adult mortality as well as maturation bias were associated with ASR (Table 1B & S2, Fig. 2C), meaning that male-biased mortalities (where more males than females die) and male-biased maturation times (where males mature later than females) were associated with female-skewed ASRs. The effect sizes were moderately strong for all three demographic predictors, whereas it was small for birth sex ratio (Table 1B). None of these results changed qualitatively when outlier data points were excluded from the analyses (Table S3A).

To assess whether these relationships were consistent between XY and ZW species, we tested the two-way interactions of each demographic variable with GSD type (Table S4, Fig. S1). The slope of the relationships between ASR and sex differences in both juvenile and adult mortality differed between XY and ZW species, as shown by the statistically significant interaction terms (GSD × juvenile mortality bias: F_1;102_ = 4.3, p = 0.041; GSD × adult mortality bias: F_1;234_ = 4.6, p = 0.034). Specifically, these effects were moderately strong in ZW species but weak in XY species (Table S4A, Fig. S1), although this difference was no longer statistically significant for adult mortality bias when outliers were excluded (Table S3B). The relationships of ASR with birth sex ratio and maturation bias did not differ between XY and ZW species (GSD × birth sex ratio: F_1;108_ = 0.305, p = 0.582; GSD × maturation bias: F_1;353_ = 2.056, p = 0.153; Table S4A, Fig. S1), and these results remained unchanged when outlier data points were excluded (Table S3B).

### Phylogenetic path analysis

The best supported model included all identified bivariate relationships plus the direct link between GSD and ASR (Model 1.b) and yielded acceptable fit with the data (Fisher’s C-statistic, p = 0.074; Fig. 3, Table 2A) according to the approach proposed by Santos[32]. AICc values indicated a substantially stronger support for this model than for any alternative (ΔAICc ≥ 3.8 in all cases, Table 2A). Note that birth sex ratio was excluded from all path models because it was consistently unrelated to both GSD and ASR in bivariate analyses (Table 1) and it was available only for a subset of species, which would restrict the analyses to a too low sample size for an informative path analysis (see Methods for details). These results were consistent with the conclusions of pairwise comparisons between nested models using likelihood ratio tests: Model 1.b outperformed Model 1.a (χ^2^ = 3.9, df = 1, p = 0.049), justifying the inclusion of a direct link between GSD and ASR (Table 2, Fig. S2). Furthermore, the more complex models that included additional effect(s) of GSD on demographic traits (Models 2-4.b) did not provide a better fit (χ^2^ < 0.1, df = 1, p > 0.9 for all comparisons; ΔAICc > 8), supporting Model 1.b. According to this most supported model, sex bias in both adult and juvenile mortality as well as in maturation time all had negative effects of similar magnitude on ASR, while the type of GSD had weaker effects on the extent of male-biased adult mortality and ASR (Table S5, Fig. 3).

**Fig. 3.**
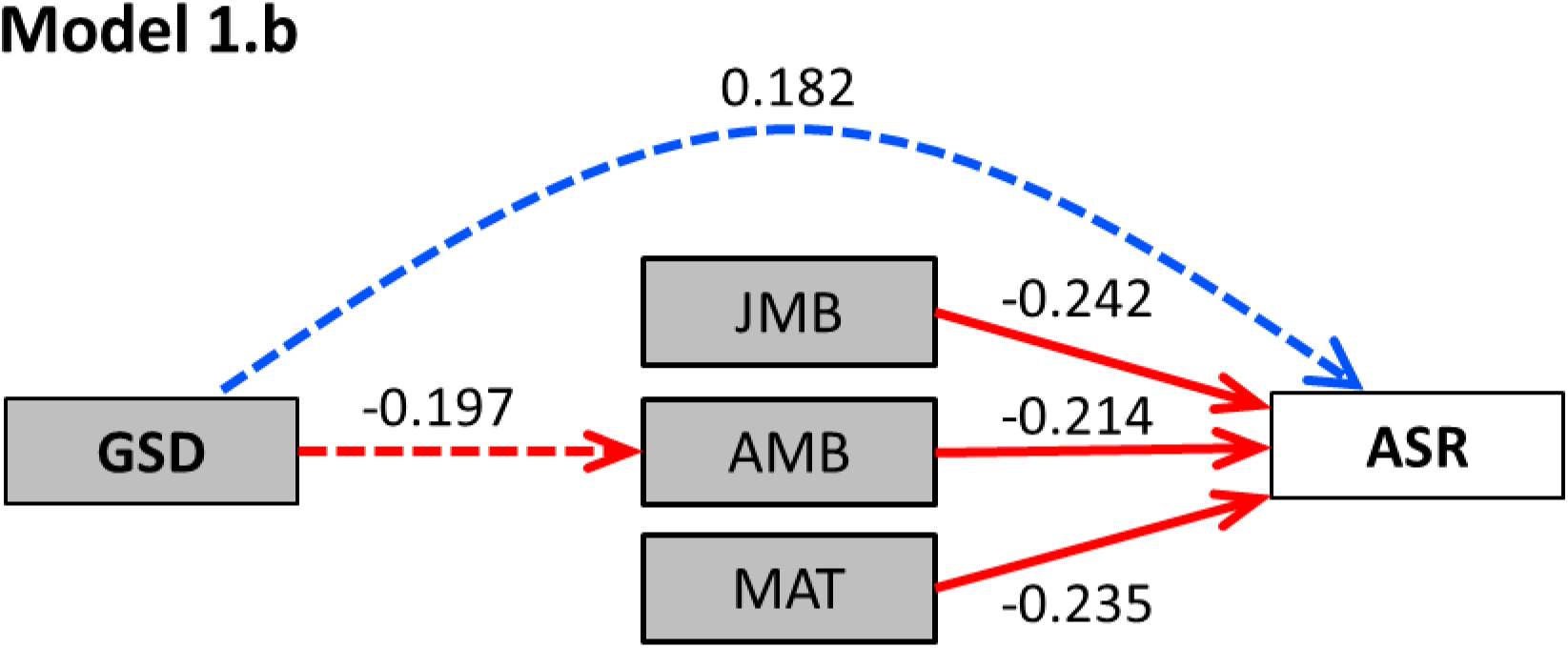
Visual interpretation of the most supported model in path analysis, (Model 1b). GSD: genetic sex determination, JMB and AMB: juvenile and adult mortality bias respectively, MAT: maturation bias, ASR: adult sex ratio. Positive and negative relationships are shown as blue and red arrows, respectively, and numbers are the standardised path coefficients (solid lines: statistically significant relationships, dashed lines: statistically non-significant relationships). See results of phylogenetic path analysis in Table 2 and Table S5, and the model set in Fig. S2.

**Table 2.**
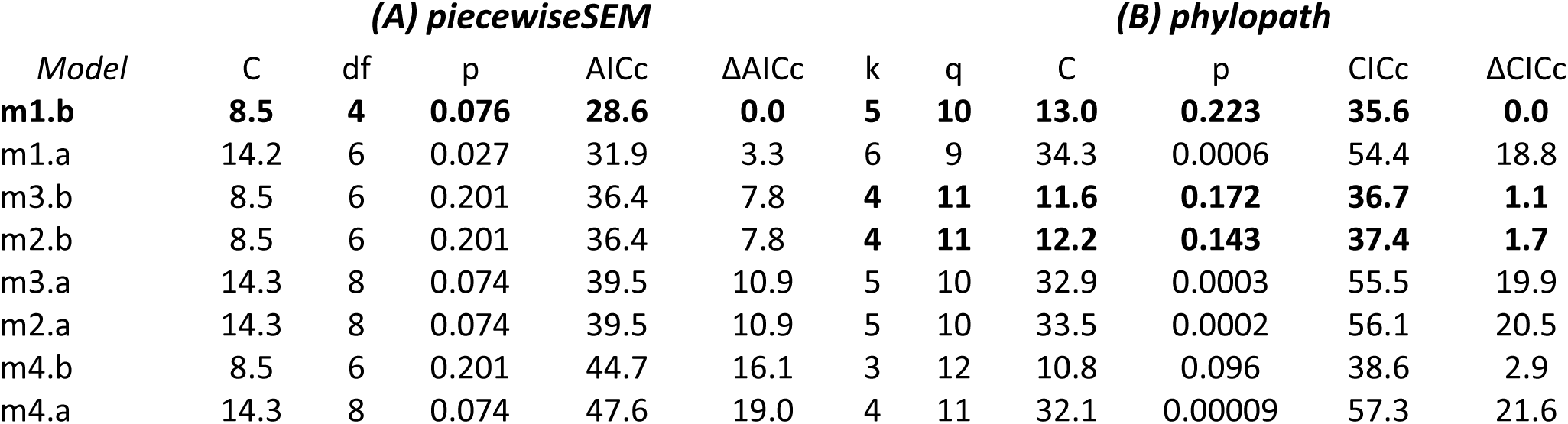
Comparison of phylogenetic path models produced **(A)** by the approach of Santos[32] using the implementation of ‘piecewiseSEM’, and **(B)** by the approach of von Hardenberg & Gonzalez-Voyer [33] using the implementation of ‘phylopath’. For each model, the table shows the number of independence claims (k), the number of parameters (q), Fisher’s C-statistic (C) for model fit and its associated p-value. AICc and CICc are the Akaike and C-statistic information criterion, respectively, corrected for small sample sizes. ΔAICc (and ΔCICc) indicates the difference in AICc (or CICc) values between the most supported model (lowest AICc or CICc, model m1.b) and the focal models. ΔAICc (and ΔCICc) > 2 indicates substantially higher support for the best model than for the other models, whereas models with ΔAICc (and ΔCICc) < 2 have similar support (highlighted in bold type). The analyses include n=95 tetrapod species with data available for all for variables, excluding birth sex ratio. The model set and path coefficients of the most supported models are given in Fig. S2 and Table S5.

These results were largely confirmed by a second set of path analyses using an alternative approach for model fitting developed by von Hardenberg and Gonzalez-Voyer[33]. The latter analyses confirmed that Model 1.b satisfactorily fits the data (Fisher’s C-statistic, p = 0.223), and model comparisons showed that this model had the lowest CICc value (Table 2B). Although two other models (Models 2.b and 3.b) had comparable support (ΔCICc < 2; Table 2B), Model 1.b was favoured as the least complex one. Results for Model 1.b corroborated that the GSD had statistically significant effects on both adult mortality bias and ASR, and that sex bias in both adult and juvenile mortality as well as in maturation time negatively influenced ASR (Fig. S2B). The two other models with ΔCICc < 2 included an additional indirect link between GSD and ASR through either male-biased juvenile mortality or maturation time, respectively (Fig. S2A). All three models showed moderately strong effects of GSD type on ASR and adult mortality bias, while the other effect sizes were smaller (Fig. S2B).

## Discussion

Our study revealed four main findings. First, we corroborated the existence of an association between the type of genetic sex-determination system and ASR using an augmented dataset and wider taxonomic coverage, i.e. that ZW species have more male-skewed ASRs than XY species (that are typically female-skewed). Second, birth sex ratios are unlikely to drive ASR variation in tetrapods. This supports and generalizes earlier studies in which no relationship was detected in birds[1,3,26,34] and other tetrapods either between hatching sex ratio and ASR, or between hatching sex ratio and type of sex chromosomes[1]. However, skewed birth sex ratios might be more important in shaping ASR in species with temperature-dependent sex determination, where highly skewed birth sex ratios are more frequent and among-species variance in birth sex ratio is much higher than in GSD species[35].

Third, our results reveal that the type of sex chromosome systems influences ASR through demographic processes. Specifically, we identified sex-biased adult mortality as a key demographic pathway linking the type of GSD to ASR in tetrapods. This new finding is based on three results: (1) the sex bias in adult mortality differed between XY and ZW species, (2) it was associated with ASR, and (3) phylogenetic path analyses supported the model that included the route between GSD and ASR *via* sex differences in adult mortality. This finding improves our understanding emerging from recent comparative studies that either showed an association between the type of GSD and lifespan or sex-specific aging rates[18,19] or reported a relationship between adult mortality bias and ASR in specific taxa[1,21,26,29]. Our study goes one step further from these previous studies by demonstrating and statistically supporting the full pathway from GSD through mortality bias to ASR skew across tetrapods, although the underlying genetic mechanisms of how sex chromosomes contribute to sex-specific mortality (e.g. through the unguarded sex chromosome or toxic Y mechanisms; see introduction) and thus the biased ASRs have still to be uncovered.

Additionally, our analyses showed that ASR variation across species was related to sex differences in juvenile mortality and maturation time, generalizing earlier findings that highlighted the importance of these sex differences in shaping ASR in birds[15,34]. Our study is the first to corroborate the effects of all of these demographic predictors of ASR (i.e. sex-biased juvenile and adult mortality and sex-biased maturation time) using a single comparative dataset. The effect sizes estimated both in the bivariate (Table 1) and path (Fig. 3) models indicated that the contribution of these demographic patterns to ASR variation are remarkably similar.

Our results provided no conclusive evidence that the type of GSD would influence ASR through sex differences either in juvenile mortality or in maturation time. While two of the supported path models included the path either through juvenile mortality bias (Model 2.b) or through maturation bias (Model 3.b), these latter results were not fully confirmed by either the bivariate models or by an alternative path analysis approach. These inconsistencies can be explained, at least in part, by reduced statistical power in some of these analyses. Specifically, the variation in the type of sex chromosome systems in the subsets of taxa used in these analyses may be limited (Figure S3), especially for juvenile mortality bias. Therefore, more research is needed to clarify whether the genetic sex-determination system has a role in generating sex differences in juvenile mortality and maturation time. The most promising way forward is to include more taxa from clades with frequent changes of the sex chromosome system, for example some lineages of amphibians and reptiles.

Besides the indirect links between GSD and ASR mediated through sex-biased adult mortality, all of our supported path models included a direct link between GSD and ASR. This can represent additional processes that were not tested in our study. For example, such a mechanism that may potentially link GSD type to ASR is variation in dispersal ability and in the magnitude of sex differences in natal or breeding dispersal. For instance, flying ability of most birds and high movement capacity of large mammals can make these groups more vagile and thus more sensitive to ASR sampling bias and/or mortality sex-bias resulting from sex-biased dispersal than other vertebrate clades[36]. Additionally, sex-biased dispersal may differ between sex-determination systems: for example, birds (ZW species) typically show female-biased dispersal, whereas mammals (XY species) show male-biased dispersal[37,38]. These sex-specific movements can influence ASR by generating a local shortage and/or higher mortality in the sex dispersing farther[2,39,40]. However, the local effects of sex-biased dispersal on ASR can be balanced out at the metapopulation level[26]. Studies so far yielded inconsistent results for the relationships between GSD, sex-biased dispersal, sex-biased mortality and ASR. Pipoly et al.[22] did not detect any difference in the magnitude of sex-biased dispersal between XY and ZW reptiles, as dispersal tended to be male-biased regardless of GSD type. The same study did not reveal any relationship between sex-biased dispersal and ASR in birds, but detected an association across tetrapods (although the latter result was inconsistent between analyses; Supplementary Material 1 in [22]). In contrast, Végvári et al. [40] found that sex-biased dispersal was associated with ASR variation but not with sex-specific mortality in birds. In sum, the available evidence is inconclusive, hence further studies would be valuable for clarifying whether sex-biased dispersal is associated with GSD and skewed ASRs.

Some of our results suggested that ASR may be more strongly associated with sex differences in adult and especially juvenile mortality in ZW species than in XY species. The reason for these differences is not clear. Theoretically, ZW systems may facilitate sex-biased mortality because sexual selection for secondary sexual characters may be more effective in ZW than in XY systems[41]. Although this may promote divergence in some demographic and life-history traits between the sexes in ZW taxa, it is unexplored whether and how this could generate stronger and/or more consistent effects on ASR than those observed in XY species. Alternatively, it is possible that the hemizygous chromosome is degenerated to a different extent in the XY and ZW systems, lending different strength to processes like the toxic Y effect for driving sex-specific life histories and population dynamics. Although the current hypotheses on the lifespan consequences of the sex chromosomes are mostly formulated for XY systems, such as the toxic Y hypothesis or the putative disappearance of the Y chromosome among cells throughout life[27], similar effects may operate in ZW systems and might be exacerbated by accumulation of retroelements[42]. Future studies should explore these possibilities, for example by taking into account the differences between the two sex chromosomes in size and gene content.

Taken together, our study consistently support that sex chromosomes can impact, via demographic pathways, the relative number of males and females. Specifically, the results infer the difference in ASR between species with XY/XX and ZZ/ZW sex-determination systems, and demonstrates that sex-biased adult mortality plays a key role in generating this difference. This conclusion, based on the largest and most diversified dataset of tetrapod species ever analysed to date, brings together the mosaic of previous results obtained across various taxon-specific analyses showing that, on one hand, the heterogametic sex may be more exposed to some deleterious effects than the homogametic sex[27,43], resulting in shorter lifespan[18,20,27] or faster ageing rate[19], while on the other hand sex differences in life histories may drive ASR biases.

Further studies are needed to uncover the mechanisms of how sex chromosomes can contribute to the differential mortality of males and females across tetrapods, which could have important implications for both population demography and reproductive trait evolution.

## Materials and Methods

### Data collection

Sex determination information was collected from the Tree of Sex database[44], with additional updates from recent primary publications (Supplementary Data). In this study we only included species that have genetic sex-determination systems (GSD): either XY/XX or ZZ/ZW sex-determination systems (with either homomorphic or heteromorphic sex chromosomes), or variants of these systems with demonstrated male or female heterogamety[44]. We did not include species with mixed sex determination, i.e. where both temperature and sex-chromosome effects simultaneously operate during sex determination[45,46]. All birds were assigned to ZZ/ZW, and all mammals to XY/XX sex determination, respectively[47]. For amphibians, only species with exactly known GSD type were included, as there can be differences even between closely related species or between populations within a given species[48]. For reptiles, we included all species for which GSD was known at least either at the genus or family level and the type of GSD was not variable within the genus or family (see [22] for details). If a species (or its genus or family) for which we had demographic information did not occur in the Tree of Sex database, we searched for its GSD in primary literature. We did not identify reliable GSD data for seven amphibian species for which we had demographic information, so these species were excluded from the analyses of GSD effects on demographic variables, but were included in the analyses of demographic trait effects on ASR.

To illustrate evolutionary changes in the sex chromosome systems in the studied clades, we mapped the transitions between sex determination systems using the ‘make.simmap’ function in the ‘phytools’ package[49] in R[50] with 100 simulations. We used two different models: the ‘equal rates’ model (ER), where only equal transition rates are allowed between the character states; and the ‘all rates different’ model (ARD), where different transition rates are allowed between the states, and compared the two models according to their AIC (Akaike Information Criterion) value. Transition reconstruction resulted on average 16.8 transitions for ARD and 18.17 for ER model between XY and ZW systems in our dataset, with an average 1.04 (ARD) or 7.01 (ER) transitions from XY to ZW systems, and an average 15.76 (ARD) or 11.09 (ER) transitions from ZW to XY systems. ARD had lower AIC value than ER (ΔAIC=3.308), hence we present the results of the ARD analysis (Fig. S3). We successfully utilized this variation in sex chromosome systems among tetrapods in our earlier studies as well. For example, Pipoly et al.[22] showed that the sex chromosome system is associated with ASR both in tetrapods as a whole and in its taxonomic subsets (amphibians and reptiles). By using a subset of 81 reptiles, Pipoly et al.[13] demonstrated the association between sex chromosome systems and the occurrence of multiple paternity. Finally, Bókony et al.[35] found differences between genetic vs. environmental sex determination systems in various demographic traits from a dataset including information on 181 reptiles. These results show that tetrapods are a suitable model system to study associations between the type of sex chromosome systems and various demographic and reproductive traits.

ASR estimates (expressed as the proportion of males in the adult population[51,52]) were taken from Pipoly et al.[22] for 344 tetrapod species living in the wild. This initial dataset was augmented with ASR data for additional amphibian, reptile and mammal species, taken from existing comparative datasets (reptiles[35], mammals[17,53]), or from primary literature searched in Web of Science or in Google Scholar (updated until June 2020; Supplementary Data). We assessed the quality of the available data records and used only those ASR estimates that were based (1) on an accurate sexing method (i.e. molecular sexing or sexual organ identification) and (2) on a sampling method for male and female numbers that was presumably not sex-biased (see further details on data quality and repeatability of ASR estimates in refs [14,17,22,35]). The final dataset includes 453 species with ASR estimates (49 amphibians, 93 reptiles, 187 birds, and 124 mammals).

We then collected information on further demographic traits from populations in the wild. First, we collected birth sex ratio data (BSR; the proportion of males at birth or hatching among the neonates in the population). We focused on the species listed in our ASR dataset and searched in Web of Science and Google Scholar using the search terms *‘(birth OR secondary) sex ratio*’* and the taxonomic name of the species. We evaluated these studies by similar criteria as the ASR studies (see above), and we only kept reliable estimates. We found reliable BSR data for 2 amphibian, 28 reptile, 55 bird and 27 mammal species (for which we also had ASR information). We included BSR estimates where the whole clutches/litters were sampled soon after hatching/birth, but sexing techniques could be variable, e.g. molecular or based on sexual organs. Wilson and Hardy[51] showed that analysing sex ratios as a proportion is reliable when sex ratios are estimated from samples of ≥ 10 individuals and the dataset has ≥ 50 sex ratio estimates, which met requirements in our analyses for both ASR and BSR.

Second, we collected data on annual juvenile mortality rates and annual adult mortality rates for males and females separately. We used data either from existing comparative datasets (birds[2]; mammals[20]), or from primary publications (Supplementary Data). In the latter case we searched the literature through the search engines Web of Science and Google Scholar, using the search terms *‘male AND female AND (mortality OR survival)’* together with the taxonomic names of the species for which ASR estimates were compiled. We categorised mortality estimates as either juvenile or adult mortality in the same way as these were provided in the original sources, i.e. as juvenile mortality when the source stated it was estimated for juveniles, immatures or subadults, and as adult mortality when the source stated that it was calculated for adults or mature individuals. Overall, juvenile mortality corresponded to the mean annual mortality prior to the age at first reproduction, whereas adult mortality corresponded to annual mortality from the age at first reproduction onwards. Mean weighted age-specific mortalities were used when fully age-dependent estimates of adult survival were available. We calculated the sex difference in annual juvenile and adult mortality rates (juvenile mortality bias and adult mortality bias, respectively) from sex-specific estimates as log_10_(male annual mortality rate / female annual mortality rate)[17,54,55], leading to measure the magnitude of the sex-biased mortality. Thus, positive values of mortality bias mean higher male than female mortality, whereas negative values mean higher female than male mortality. We were able to estimate juvenile mortality bias for 108 tetrapod species (3 amphibians, 17 reptiles, 52 birds and 36 mammals), and adult mortality bias for 244 species (23 amphibians, 41 reptiles, 128 birds and 52 mammals).

Third, we collected data on the sex-specific age of sexual maturity (in days). We used either existing datasets (for anurans [56], for reptiles [35], for birds [15], and for reptiles and mammals [57]), or searched the literature using the search terms *‘male AND female AND age AND (matur* OR first reproduction)’* together with taxonomic names. We quantified the sex differences in maturation time (referred to in this paper as maturation bias) as log_10_(male maturation age/ female maturation age). Positive values mean that males mature later than females, whereas negative values mean that females mature later than males. We were able to estimate maturation bias for 366 species (37 amphibians, 70 reptiles, 175 birds and 84 mammals).

Most data on sex ratios, mortality and maturation were collected blindly with respect to the hypotheses we test in the current analyses because these data were originally collected for the purposes of other comparative studies. Data on different demographic traits for the same species were often available only from different populations or studies. For all demographic traits, we used the mean value for each species whenever reliable data were available from more than one population or study.

### Statistical analyses

All analyses were run in R (version 4.0.0; R Core Team 2020)^52^. We standardised all numeric variables by centring to the mean and dividing by standard deviation, and we used these values for all analyses in main text. We built phylogenetic generalised least squares (PGLS) models to conduct bivariate analyses using the ‘pgls’ function of the ‘caper’[58] R package. To control for the phylogenetic relatedness among taxa, we used the composite phylogeny applied in [22] with the addition of new species according to a family-level reptile phylogeny[48] and other recent phylogenies (Squamata[59,60], Testudines[61], birds[62], mammals[63,64]). Since composite phylogenies do not have true branch lengths, we used Nee’s method to generate branch lengths using Mesquite software[65]. In each model we used the phylogenetic signal (i.e. Pagel’s λ[66]) as estimated by the maximum-likelihood method for that model.

Using all species with available data, we first applied two sets of bivariate PGLS models. In the first set of analyses, we tested whether each of the examined demographic traits differed between the two types of GSD (i.e. XY versus ZW species). In these models, the type of GSD was the predictor variable, and one of the four demographic trait (birth sex ratio, juvenile mortality bias, adult mortality bias and maturation bias) was the dependent variable. In the second set of analyses, we assessed whether demographic traits predict ASR. In these models, ASR was the response variable and one of the four demographic traits was the predictor variable. To check whether a non-linear relationship would fit the data better than a linear model, we re-ran this second set of models with adding the quadratic term of the predictor. However, these latter models never provided a better fit than the ones without the quadratic term (Table S6).

We also tested whether the relationships of ASR with the demographic predictors differed between GSD types. For this analysis, we built PGLS models using the ‘gls’ function of the ‘ape’[67] R package, with the corPagel correlation structure accounting for phylogenetic relationships among species. In these models, ASR was the response variable, and one of the demographic traits, GSD type, and demographic trait × GSD type interaction were included as predictors. We used type-3 ANOVA tables (R function ‘anova.gls’ with the argument type=‘marginal’) to test whether the interaction was statistically significant in each of the above models. Then, for each interaction model we tested whether the slope of the relationship between ASR and the demographic variable differed from zero in either of the two GSD types (XY or ZW species) using the ‘emtrends’ function of package ‘emmeans’[68].

The validity of the models was checked by inspection of residual diagnostic plots to ensure that all statistical assumptions were met. We also checked the sensitivity of the results to the influence of outlier data points, which were identified from the standardised predictor variables as being more than three standard deviations distance from the mean (i.e. have high leverage). Outliers were identified separately for each group (XY or ZW) within each model. We repeated all PGLS analyses with demographic predictors after excluding these potentially highly influential data points (see Table S3 for number and identity of species excluded). Throughout the paper, we present figures using the untransformed data points (i.e. non-standardised values) to facilitate interpretation. We repeated the bivariate analyses using non-standardised variables (Table S2) to allow for the interpretation of the effect sizes on the original data scales. For illustration purposes, we added regression lines to the figures calculated from PGLS models using non-standardised data, and plotted the uncertainty of the slopes using the ‘evomap’[69] package in R. In tables of the main text, we present the results with the transformed (i.e. standardised) variables.

We used phylogenetically controlled confirmatory path analyses to investigate the fit of models representing different sets of pathways linking GSD and ASR (Fig. S2A). First, we constructed a path model that included links (1) between GSD and adult mortality bias, and (2) between ASR and three demographic traits: juvenile and adult mortality bias, and maturation bias (Model 1.a). This model aimed to simultaneously estimate the relationships that were detected in the bivariate PGLS analyses (see Results and Table 1), and also corresponded to the existing knowledge on the effect of GSD on sex-specific lifespan[18,19] and the association of ASR with sex-biased mortality and maturation[15,26,30]. We did not include the path leading from BSR to ASR because it was consistently not supported both by our current results (see Table 1) and earlier analyses[2]. Furthermore, BSR was available only for a relatively small subset of species, thus by excluding this variable we were able to double the sample size of the path models (n= 43 and 95 species, with BSR included or excluded, respectively), resulting in a more suitable sample size for path analyses[70]. Then, we created an extended model by adding the direct link between GSD and ASR (Model 1.b). Finally, we created three further expanded versions of both Models 1.a and 1.b, each of which contained additional paths between GSD and (1) juvenile mortality bias, (2) maturation bias or (3) both juvenile mortality and maturation bias (Models 2-4.a and 2-4.b). By these latter models, we tested whether the fit of path models can be improved by including relationships that were not supported in the bivariate PGLS analyses. Note that Model 1.a is nested in Model 1.b, and both of these models are nested in Models 2-4a and 2-4b, respectively, facilitating model comparison (see below). We used those species for path analyses for which we had data for all included variables (i.e. ASR, GSD, juvenile and adult mortality bias and maturation bias; n= 95 species).

We used two methods to fit the path models to the data. Firstly, we followed the approach proposed by Santos[32] that applies phylogenetic transformation on the data before the path analysis to control for the effects of phylogenetic relatedness among species. For this purpose, Pagel’s λ was estimated separately for each variable by maximum likelihood, and this variable-specific λ value was used to re-scale the phylogenetic tree. Then we used this transformed tree to calculate phylogenetically independent contrasts for the variable using the ‘pic’ function of the R package ‘ape’[67]. We repeated this process for each variable, and the resulting phylogenetically-transformed variables were used for fitting the path models by the R package ‘piecewiseSEM’[71], which implements the d-separation method[72]. This approach uses Fisher’s C-statistic to test the goodness of fit: the model is rejected (i.e. it does not provide a satisfactory fit to the data) if the C-statistic is statistically significant, and conversely a high p-value means satisfactory fit[71]. Model support was compared between different path models by their AICc (Akaike Information Criterion corrected for low sample sizes) values. Since information-theoretic model comparisons (e.g. those based on AICc) can be unreliable for path models[70,73] (see also Appendix 3 in ref[17] for an example in the context of comparative analyses), we also conducted pairwise model comparisons using likelihood ratio tests (LRT). LRT is straightforward to conduct for pairs of nested models, where a statistically significant difference supports the model with higher likelihood (better fit)[74]. For LRTs, we conducted the model fitting using the ‘lavaan’ R package[75] and used the ‘anova’ function to compare pairs of nested models.

Secondly, to test the robustness of the results, we repeated the path analysis using the method developed by von Hardenberg & Gonzalez-Voyer[33]. Unlike Santos’ method[32], in this latter approach a single value of Pagel’s λ is estimated for each pair of traits using PGLS models, rather than a value of λ for each variable. Model fitting and model comparisons were conducted using the ‘phylopath’ package[76]. Model fit was evaluated using the d-separation method[72] and compared between path models by their C-statistic Information Criterion (CICc) corrected for small sample sizes [33]. CIC is a modified version of AIC developed for d-separation path analysis framework[77], and is used in the same way as AIC to compare model fit. Confidence intervals for the standardised path coefficients were estimated by bootstrapping, using the ‘boot’ function with 500 iterations.

## Acknowledgments

We thank D. Gigler, G. Mészáros, G. Milne, H. Naylor and E. Sebestyén for help in data collection. This project has received funding from the HUN-REN Hungarian Research Network.

## Data and materials availability

Our dataset used in the analyses (Supplementary Data) is publicly available after acceptance and publication of the study. All data are available in the main text or the supplementary materials. All analyses were done in the R free statistical software environment.

---

**Fig. S1.**
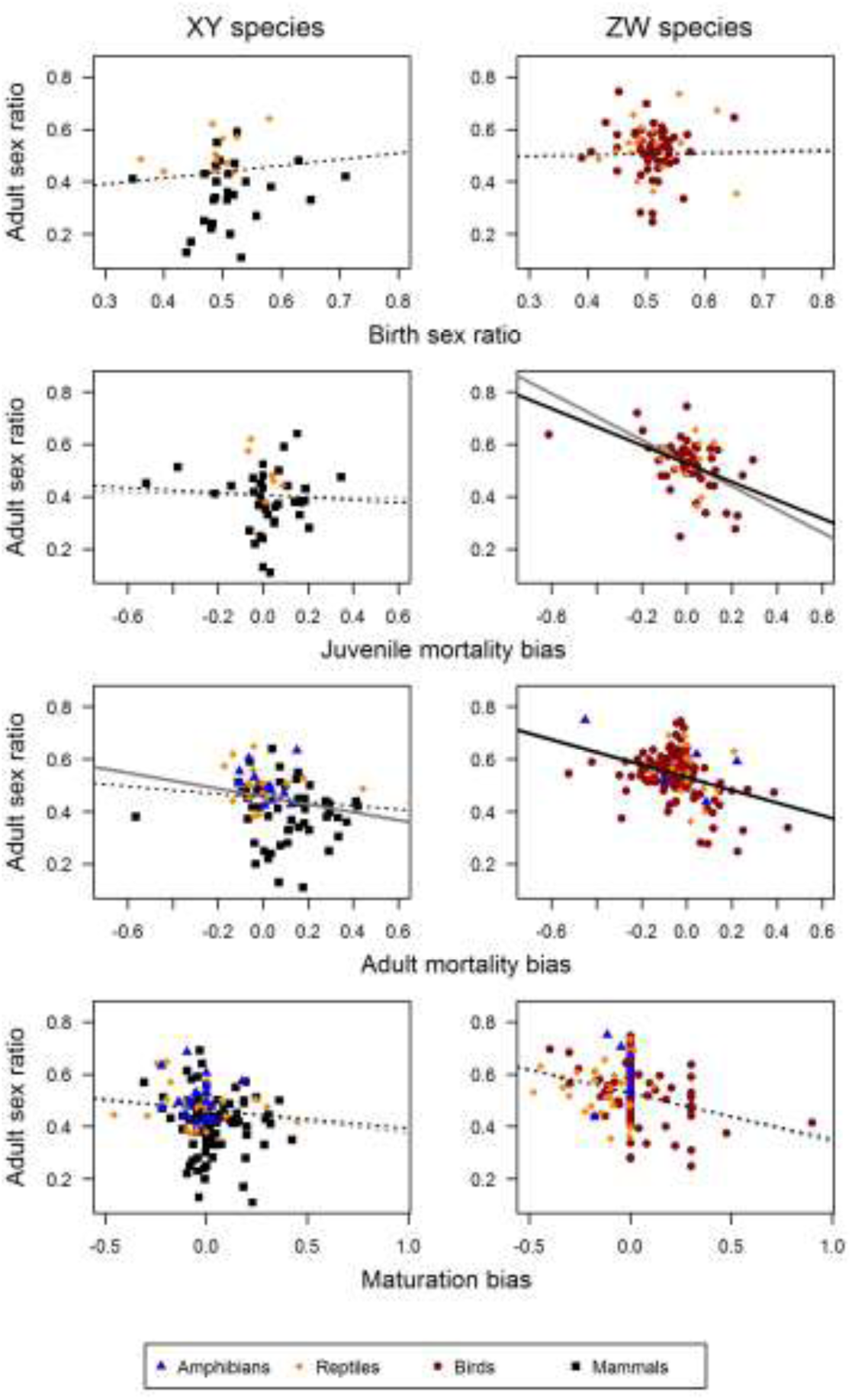
Relationships of ASR with demographic traits (i.e. birth sex ratio, juvenile mortality bias, adult mortality bias, and maturation bias) in XY and ZW species separately. On the axes, >0.5 sex ratio values mean male-skewed sex ratios and <0.5 values mean female-skewed sex ratios, while positive bias values mean higher male than female mortalities/maturation times, whereas negative bias values mean higher female than male mortalities/maturation times. Regression lines in the figures are calculated from PGLS models using non-standardised data. Statistically significant relationships are illustrated with continuous lines, whereas not statistically significant relationships are shown with dashed regression lines (see Table S3). For each panel, black regression lines are from models with all data points, and grey regression lines are from models without outlier data points (Table S2).

**Fig. S2.**
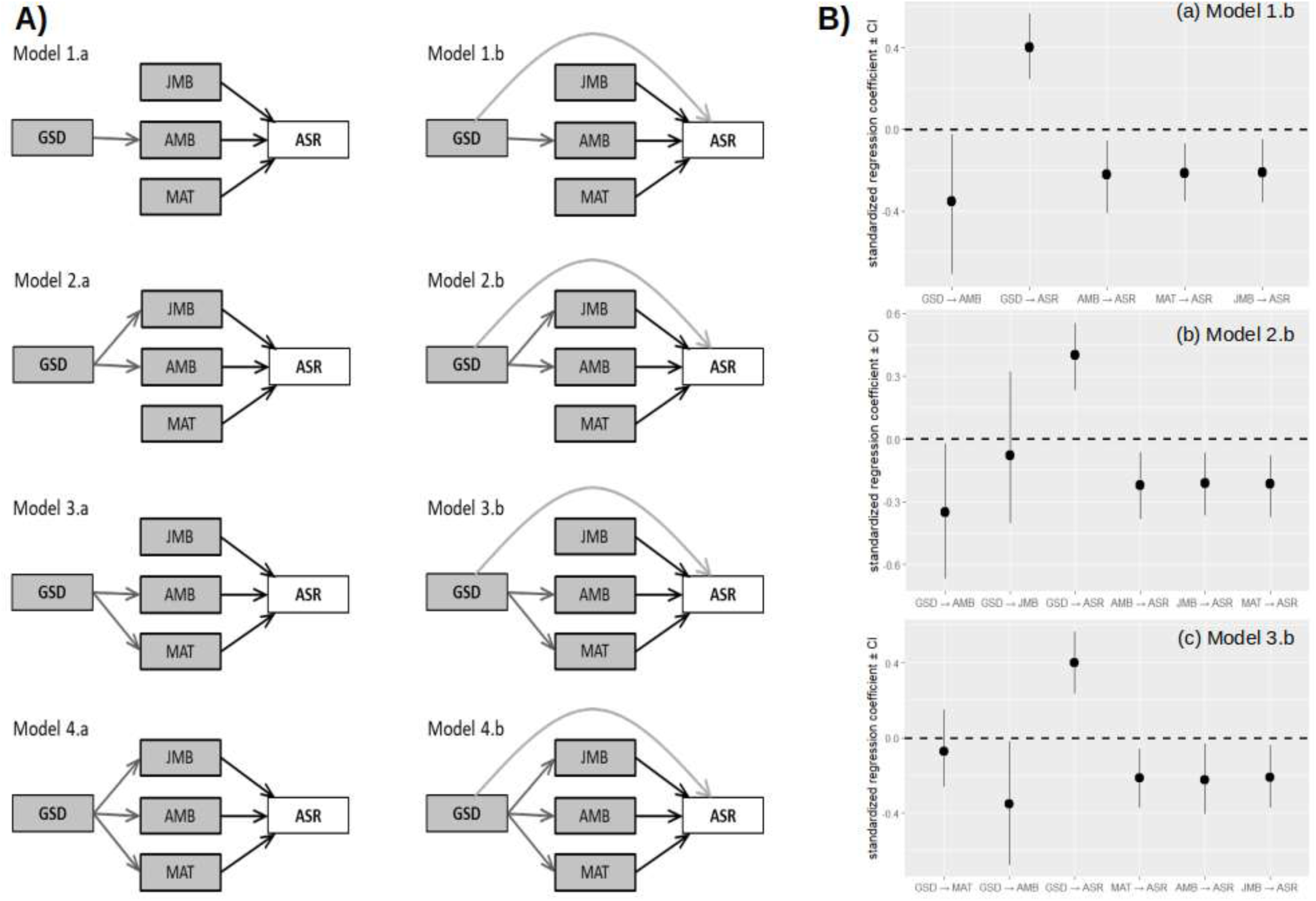
Visual representation of (A) our model set for path analysis, showing all models we tested, and (B) the standardised path coefficients and associated confidence intervals for the most supported models as inferred by analyses using R packge ‘phylopath’. In A), the tested combinations of potentially important direct (light grey arrows) and indirect (dark grey arrows) links from genetic sex determination to adult sex ratio are presented. Influence of the three demographic traits on adult sex ratio (black arrows) were included in all models. In B), panels present the standardized path coefficients inferred by each supported model. Confidence intervals were estimated by bootstrapping, using the ‘boot’ function with 500 iterations. Statistics for the model fit and comparisons are presented in Table 2B in the main text. GSD: genetic sex determination, JMB and AMB: juvenile and adult mortality bias, MAT: maturation bias, ASR: adult sex ratio.

**Fig. S3.**
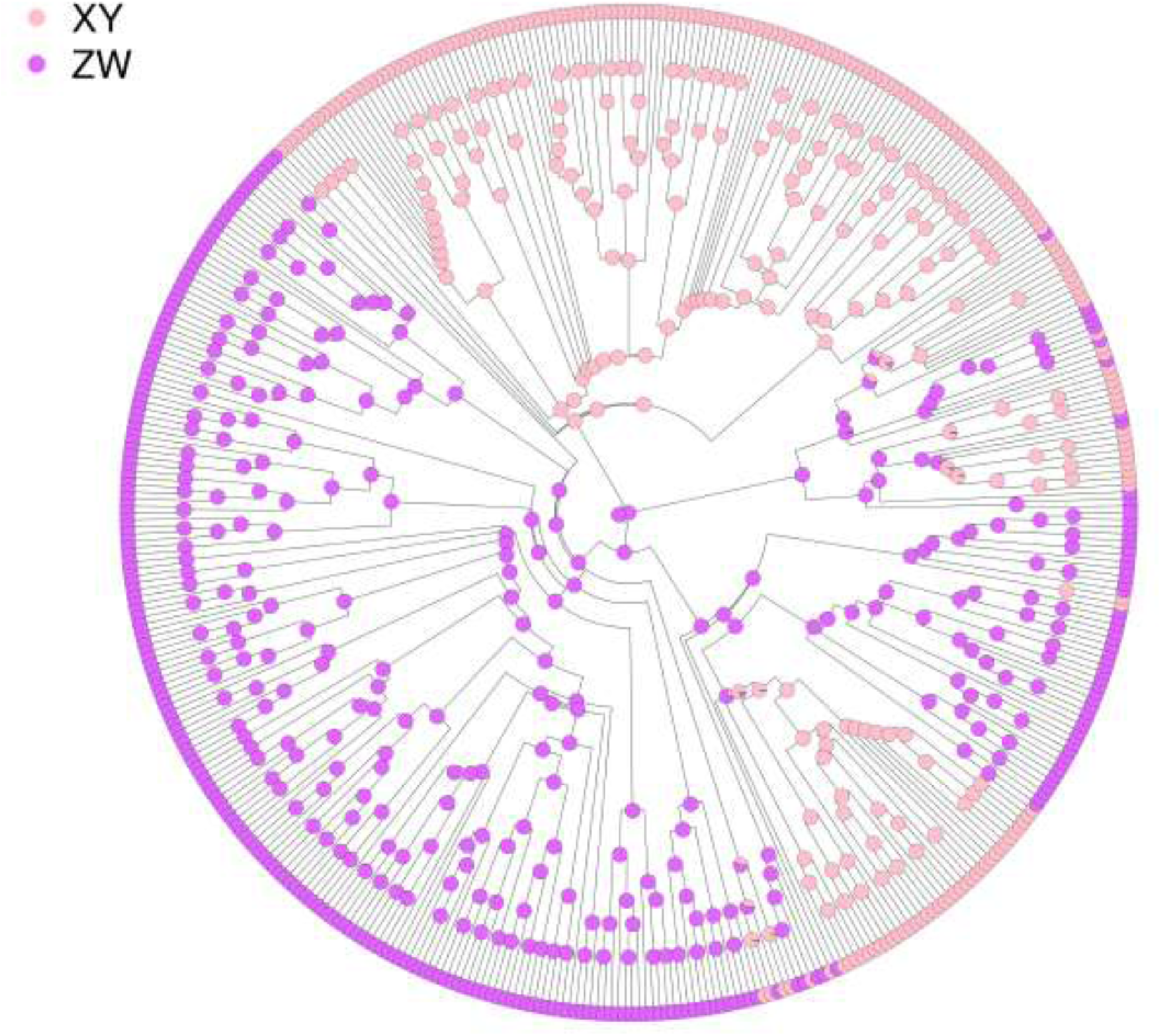
Result of transition reconstruction between XY and ZW sex determination systems in tetrapod clades included in the study (n=446 species with known sex determination system). We used the ‘make.simmap’ function in the ‘phytools’ package^52^ in R^53^ with 100 simulations. Result of the “all rates different” (ARD) model type is shown, where different transition rates are allowed between the states, because this model type was supported most by its AIC value. Pie charts on each node indicate the likelihood of the two SD system types at a given node, and the central node represents the estimated common ancestor.

**Table S1.**
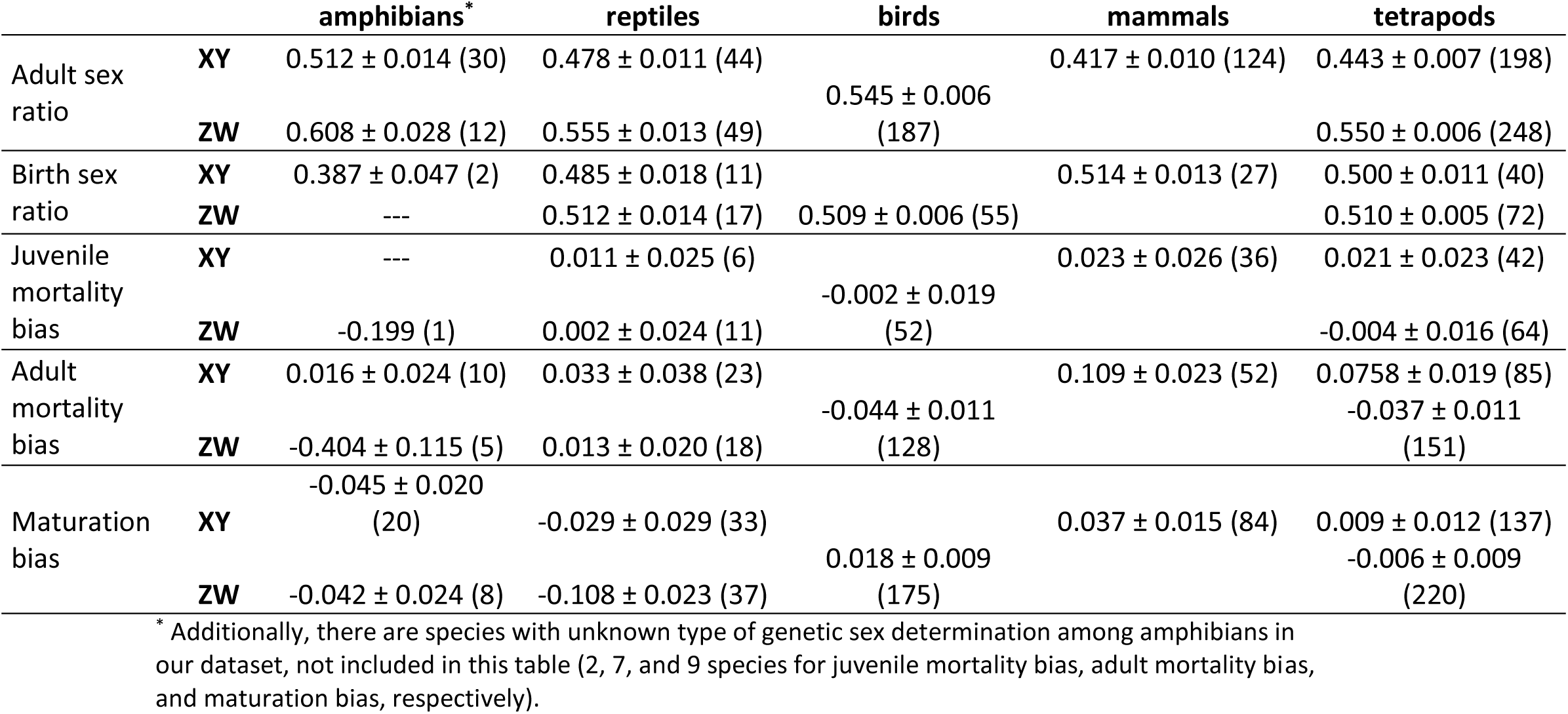
Descriptive statistics for the demographic traits in XY and ZW species in separate taxonomic groups and overall in tetrapods. Mean ± SE values estimated from non-standardised data are presented, followed by number of species in brackets. For statistical comparisons among groups, see Table S3.

**Table S2.** Results of phylogenetic generalised least squares models with the raw, non-standardised values (see Methods for details) A) between each examined demographic variable separately and GSD type across tetrapods, B) between ASR and each examined demographic variable across tetrapods, Table shows the results of bivariate PGLS models. In section A, GSD type is the predictor, and demographic variables are the response variables, separately each. In section B), the response variable is the ASR in all models. **λ** is the phylogenetic signal (Pagel’s lambda), N shows the number of species. Statistically significant (i.e. p<0.05) relationships are highlighted in bold.

| A) | $b \pm SE$ | t | p | $\lambda$ | N |
| --- | --- | --- | --- | --- | --- |
| Adult sex ratio ~ GSD | $0.086 \pm 0.018$ | 4.715 | **< 0.001** | 0.316 | 446 |
| Birth sex ratio ~ GSD | $0.010 \pm 0.011$ | 0.889 | 0.376 | 0.000 | 112 |
| Juvenile mortality bias ~ GSD | $-0.028 \pm 0.043$ | -0.639 | 0.524 | 0.194 | 106 |
| Adult mortality bias ~ GSD | $-0.081 \pm 0.038$ | -2.152 | **0.032** | 0.307 | 238 |
| Maturation bias ~ GSD | $-0.048 \pm 0.030$ | -1.595 | 0.112 | 0.356 | 357 |
| B) | $b \pm SE$ | t | p | $\lambda$ | N |
| Birth sex ratio | $0.190 \pm 0.182$ | 1.045 | 0.298 | 0.398 | 112 |
| Juvenile mortality bias | $-0.206 \pm 0.079$ | -2.615 | **0.010** | 0.338 | 108 |
| Adult mortality bias | $-0.170 \pm 0.040$ | -4.198 | **< 0.001** | 0.280 | 242 |
| Maturation bias | $-0.142 \pm 0.036$ | -3.928 | **< 0.001** | 0.380 | 364 |

In A) "b" shows the difference in ASR between XY and ZW species, with XY as the reference group. In B) "b" shows the slope of the relationship between ASR and demographic variables.

**Table S3.** Associations without the potential influential data points (see Methods for details) A) between ASR and each examined demographic trait across tetrapods, B) between ASR and the interaction of demographic traits and GSD type. Table shows the results of bivariate PGLS models (A), and results of PGLS models including the two-way interaction of each demographic trait with GSD type (B). **λ** is the phylogenetic signal (Pagel’s lambda), N shows the number of species, N_excl_ shows the number of excluded influential data points for each model. Note that the continuous variables were standardised to 0 mean and 1 SD, so the model coefficients (b) are comparable across predictors. Statistically significant (i.e. p<0.05) relationships are highlighted in bold.

| Predictor variable | $b \pm SE$ | t | p | $\lambda$ | N | $N_{\text{excl}}$ |
| --- | --- | --- | --- | --- | --- | --- |
| **A)** |  |  |  |  |  |  |
| **Birth sex ratio** |  | No influential points detected |  |  |  |  |
| **Juvenile mortality bias** | -0.304 ± 0.122 | -2.491 | **0.014** | 0.324 | 106 | 2 |
| **Adult mortality bias** | -0.351 ± 0.069 | -5.064 | **< 0.001** | 0.263 | 238 | 4 |
| **Maturation bias** | -0.183 ± 0.057 | -3.195 | **0.002** | 0.383 | 356 | 8 |
| **B)** |  |  |  |  |  |  |
| | F | df | p | $\lambda$ | N | $N_{\text{excl}}$ |
| **Birth sex ratio × GSD** | 3.385 | 1;106 | 0.069 | 0.359 | 110 | 2 |
| **Juvenile mortality bias × GSD** | 5.526 | 1;100 | **0.021** | 0.063 | 104 | 2 |
| **Adult mortality bias × GSD** | 0.807 | 1;227 | 0.369 | 0.172 | 231 | 7 |
| **Maturation bias × GSD** | 1.332 | 1;347 | 0.249 | 0.308 | 351 | 6 |

In A) "b" shows the slope of the relationship between ASR and demographic variables.
In B) and C), results of type 3 ANOVA are presented.
Excluded species for bivariate models: Birth sex ratio: no influential points detected; Juvenile mortality bias: *Loxioides\_bailleui*, *Bison\_bison*; Adult mortality bias: *Tayassu\_pecari*, *Crocutea\_crocutea*, *Ctenosaura\_melanosterna*, *Pogonocichla\_stellata*; Maturation bias: *Antidorcas marsupialis*, *Hynobius leechii*, *Cisticola\_juncidis*, *Tetrao\_urogallus*, *Ameiva\_quadrilineata*, *Anolis\_garmani*, *Apalone\_spiniifera*, *Vipera\_aspis*. Excluded species for GSD interactions: Birth sex ratio: *Megalurus\_gramineus*, *Iberolacerta\_aranica*; Juvenile mortality bias: *Loxioides\_bailleui*, *Bison\_bison*; Adult mortality bias: *Tayassu\_pecari*, *Crocutea\_crocutea*, *Ctenosaura\_melanosterna*, *Pogonocichla\_stellata*, *Circus\_cyaneus*, *Selasphorus\_platycercus*, *Bufo\_bufo*; Maturation bias: *Cisticola\_juncidis*, *Tetrao\_urogallus*, *Ameiva\_quadrilineata*, *Anolis\_garmani*, *Apalone\_spiniifera*, *Vipera\_aspis*.
Excluded species for Taxon interactions: Birth sex ratio: *Megalurus\_gramineus* (bird); Juvenile mortality bias: *Loxioides\_bailleui* (bird), *Bison\_bison* (mammal); Adult mortality bias: *Tayassu\_pecari* (mammal), *Crocutea\_crocutea* (mammal), *Ctenosaura\_melanosterna* (reptile), *Pogonocichla\_stellata* (bird), *Circus\_cyaneus* (bird), *Selasphorus\_platycercus* (bird); Maturation bias: *Hynobius leechii* (amphibian), *Cisticola\_juncidis* (bird), *Tetrao\_urogallus* (bird), *Jacana spinosa* (bird), *Ameiva\_quadrilineata* (reptile).
Note that amphibians are excluded from models of birth sex ratio and juvenile mortality bias due to lack of enough data for this group.

**Table S4.** Relationships between ASR and demographic traits separately for XY and ZW species. Continuous variables are standardised in all models. Slopes of the relationships for each group were taken from models that include interaction between demographic traits and GSD type (XY or ZW). Estimates are based on models in which continuous variables were standardised to 0 mean and 1 SD. Slopes significantly differing from zero, i.e. with 95% confidence interval (CI) excluding zero, are highlighted in bold. **λ** is the phylogenetic signal (Pagel’s lambda), and N shows the number of species in each model, and the number of species in each presented group (in italics).

| Predictor | Slope ± SE | 95% CI | $\lambda$ | N |
| --- | --- | --- | --- | --- |
| *Comparing the ASR - demography relationships between XY and ZW species* |  |  |  |  |
| **<u>Birth sex ratio</u>** |  |  | 0.352 | 112 |
| GSD type: XY | 0.121 ± 0.125 | -0.127 ; 0.370 | | 40 |
| GSD type: ZW | 0.017 ± 0.140 | -0.261 ; 0.296 | | 72 |
| **<u>Juvenile mortality bias</u>** |  |  | 0.057 | 106 |
| GSD type: XY | -0.059 ± 0.133 | -0.324 ; 0.260 | | 42 |
| GSD type: ZW | **-0.435 ± 0.123** | **-0.679 ; -0.191** |  | 64 |
| **<u>Adult mortality bias</u>** |  |  | 0.184 | 238 |
| GSD type: XY | -0.107 ± 0.080 | -0.265 ; 0.051 | | 87 |
| GSD type: ZW | **-0.353 ± 0.083** | **-0.516 ; -0.190** |  | 151 |
| **<u>Maturation bias</u>** |  |  | 0.307 | 357 |
| GSD type: XY | -0.091 ± 0.069 | -0.227 ; 0.045 | | 137 |
| GSD type: ZW | **-0.224 ± 0.062** | **-0.345 ; -0.102** |  | 220 |

**Table S5.** Estimates of non-standardised and standardised coefficients, and associated SE and p-values for paths in the most supported path model (Model 1.b) in the analysis by ‘piecewiseSEM’. SE are only estimated for non-standardised coefficients in ‘piecewiseSEM’.

| Path | Non-standardised<br>path coefficient ± SE | Standardised<br>path coefficient | Z | p |
| --- | --- | --- | --- | --- |
| GSD → ASR | 0.105 ± 0.054 | 0.182 | 1.937 | 0.056 |
| GSD → AMB | -0.153 ± 0.080 | -0.197 | - 1.922 | 0.058 |
| JMB → ASR | - 0.207 ± 0.081 | -0.242 | - 2.551 | **0.013** |
| AMB → ASR | - 0.158 ± 0.071 | -0.214 | - 2.217 | **0.029** |
| MAT → ASR | - 0.242 ± 0.094 | -0.235 | - 2.559 | **0.012** |

**Table S6.** Testing the quadratic effect of demographic predictors on ASR. ΔAIC values show the difference between the model including a quadratic term of each demographic predictor and the model without any quadratic term (i.e. positive ΔAIC value means lower AIC value for the model without the quadratic term). P-values show the result of a likelihood ratio test between the two corresponding models with and without the quadratic term, where significant (<0.05) p-value suggests better fit for models with lower AIC (i.e., without quadratic terms).

| Predictor | ΔAIC | p |
| --- | --- | --- |
| Birth sex ratio | 6.297 | 0.038 |
| Juvenile mortality bias | 6.288 | 0.038 |
| Adult mortality bias | 7.479 | 0.019 |
| Maturation bias | 8.342 | 0.012 |

